# Spatio-temporal patterns of population responses in the visual cortex under Isoflurane: from wakefulness to loss of consciousness

**DOI:** 10.1101/2020.07.20.212472

**Authors:** Shany Nivinsky Margalit, Neta Gery Golomb, Omer Tsur, Aeyal Raz, Hamutal Slovin

## Abstract

Anesthetic drugs are widely used in medicine and research to mediate loss of consciousness (LOC). Despite the vast use of anesthesia, how LOC affects cortical sensory processing and the underlying neural circuitry, is not well understood. We measured neuronal population activity in the visual cortices of awake and isoflurane anesthetized mice and compared the visually evoked responses under different levels of consciousness. We used voltage-sensitive dye imaging (VSDI) to characterize the temporal and spatial properties of cortical responses to visual stimuli over a range of states from wakefulness to deep anesthesia. VSDI enabled measuring the neuronal population responses at high spatial (meso-scale) and temporal resolution from several visual regions (V1, extrastiate-lateral (ESL) and extrastiate-medial (ESM)) simultaneously. We found that isoflurane has multiple effects on the population evoked response that augmented with anesthetic depth, where the largest changes occurred at LOC. Isoflurane reduced the response amplitude and prolonged the latency of response in all areas. In addition, the intra-areal spatial spread of the visually evoked activity decreased. During visual stimulation, intra-areal and inter-areal correlation between neuronal populations decreased with increasing doses of isoflurane. Finally, while in V1 the majority of changes occurred at higher doses of isoflurane, higher visual areas showed marked changes at lower doses of isoflurane. In conclusion, our results demonstrate a reverse hierarchy shutdown of the visual cortices regions: low-dose isoflurane diminishes the visually evoked activity in higher visual areas before lower order areas and cause a reduction in inter-areal connectivity leading to a disconnected network.

## Introduction

Despite extensive research over the past few decades the neural mechanisms mediating conscious experiences are largely unknown. Anesthetic drugs are an important research tool in this field as they enable a controllable approach for studying the transition from wakefulness to loss of consciousness (LOC). LOC can be induced by various anesthetic agents, many of them act on various molecular targets and thereby induce multiple and heterogeneous changes in neural responses (Campagna et al., 2003; Franks, 2008; Rudolph and Antkowiak, 2014). Here, we investigated the effects of isoflurane, a widely used anesthetic targeting both excitatory and inhibitory synaptic transmission (Franks, 2008). Isoflurane was previously reported to enhance inhibitory synapses by acting on GABA and glycine receptors (Hentschke et al., 2005) and also suppresses excitatory synapses through cholinergic, serotonin and glutamate/NMDA receptors (Campagna et al., 2003; Mashour, 2005). A recent study reported that inhaled anesthetics directly target the lipid membrane, thus disrupting its function (Pavel et al., 2020).

The effects of isoflurane on different types of synapses and channels generates heterogeneous responses altering neural activity in many brain regions (Liang et al., 2012a, 2015). Isoflurane was shown to affect information processing at multiple levels from local synaptic influences to global changes in functional connectivity (Franks, 2008; Hudetz and Mashour, 2016; Shushruth, 2013). Isoflurane was reported to induce activity changes in the thalamus (White and Alkire, 2003) which serves as a central sensory-motor information processing junction and was suggested to play a key role in drug-induced LOC (Bonhomme et al., 2012; Boveroux et al., 2010; Gao et al., 2017; Nallasamy and Tsao, 2011). Indeed, the thalamo-cortical pathway is suppressed under increasing levels of isoflurane (Alkire et al., 2000; Liang et al., 2012a). However, recent studies revealed that isoflurane has an even stronger suppressive effects on cortico-cortical pathways (Alkire et al., 2008; Imas et al., 2005; Raz et al., 2014; Schrouff et al., 2011; Sellers et al., 2015). Moreover, isoflurane was reported to interfere with the functional connectivity between brain areas, influencing both short and long distance connections (Liang et al., 2012a; Nallasamy and Tsao, 2011; Redinbaugh et al., 2020).

Despite the accumulating knowledge, how isoflurane affects cortical visual processing is not well understood. At the population level, isoflurane reduced the firing rates of neurons in the primary visual cortex (V1), increased their response latencies and prolonged evoked responses (Niell and Stryker, 2010; Vaiceliunaite et al., 2013). The effects on receptive fields (RF) properties of V1 neurons, is controversial: some studies showed no effect while others reported on marked changes (Aasebø et al., 2017; Andermann et al., 2011; Goltstein et al., 2015; Marshel et al., 2011; Niell and Stryker, 2010; Pack et al., 2001; Vaiceliunaite et al., 2013). In addition, most previous studies of drug induced LOC focused on a single drug dosage i.e. a single level of anesthesia and reported the effects at the single-neuron or large network level. Thus, the effects of isoflurane on spatio-temporal patterns of cortical responses to visual stimuli, under increasing depths of isoflurane anesthesia are not well understood. Voltage-sensitive dye imaging (VSDI) at the mesoscale approach, allows bridging between the single neuron and the large scale network studies (Berger et al., 2007; Civillico and Contreras, 2012; Devonshire et al., 2010; Ferezou et al., 2006; Petersen et al., 2003; Polack and Contreras, 2012). This approach enables us to address previously unresolved questions such as the local spatial spread of the visually evoked response and synchronization among closely located visual areas.

We measured neural activity in the visual cortex of the mouse and compared the visual evoked cortical responses under different levels of anesthesia consciousness. We used VSDI to characterize the temporal and spatial properties of cortical responses to visual stimuli, from wakefulness to deeply anesthetized state. The VSD signal enabled to investigate the spread of activation within each area and the synchronization within and between areas, under varying levels of isoflurane.

## Materials and methods

### Animals

We used 21 male C57BL\6JOlaHsd mice (12-15 weeks) in this study, with a total of 45 recording sessions (in 3 mice we collected data under two different isoflurane levels). All experimental and surgical procedures were carried out according to the NIH guidelines, approved by the Animal Care and Use Guidelines Committee of Bar-Ilan University and supervised by the Israeli authorities for animal experiments.

### Determination of loss of righting reflex

Loss of righting reflex (LORR) is a well-established method to assess behavioral responsiveness in rodents. It is used to determine responsiveness to various anesthetic agents in non-noxious manner, relying on loss of vestibular inputs when the rodent is on its back (Dickinson et al., 2000; Eger and MacLeod, 1995; Johannesson et al., 1984; Raz et al., 2014; Sonner et al., 2007). The isoflurane concentration in which LORR occurred was determined for each animal prior to surgery using SomnoSuite Small Animal Anesthesia System (Kent Scientific Corporation). The animal was placed in a 0.5 litter chamber and using 500 ml/min flow, we gradually increased the Isoflurane concentration, starting from 0.5% with increments of 0.1% every 5 minutes until LORR was achieved and documented.

### Surgery and staining

Craniotomy was performed under deep isoflurane anesthesia. The mouse was ventilated using a SomnoSuite Small Animal Anesthesia System face mask throughout the surgery and imaging. After skin incision, a chamber was cemented to the cranium with dental acrylic cement over the visual areas (centered at ~3 mm lateral from midline and ~2.5 mm posterior from lambda). A 5 mm diameter craniotomy was performed and the dura mater was removed in order to expose an imaging window over the visual cortices. The cortical surface was then carefully washed with artificial cerebrospinal fluid (ACSF) and prepared for staining. The chamber was filled with dye solution (RH-1691; 0.5 mg/ml of ACSF). The cortical staining lasted for 1.5-2 hr, the cortical surface was then washed with ACSF until the solution was clear. The chamber was filled with transparent agar and covered with transparent acrylic glass lid. The mouse was placed on Harvard Small Animal Physiological Monitoring System (Harvard Appartus), maintaining body temperature at 36.5-37^0^C and oxygen saturation > 90% in room air. Buprenorphin (SC, 0.05 mg/kg; an analgesic drug) was given to the animals before waking them and VSDI started at least 40 minutes after the Buprenorphin application. In some of the animals Buprenorphin was applied also at surgery onset.

### Anesthesia protocol

Imaging started in the awake state, 40-45 minutes after Isoflurane was stopped and while verifying normal body temperature and oxygen saturation > 90%. Awake state was confirmed by eye blinking, reflex re-appearance (e.g. withdrawal reflex) and whisking. During the anesthesia conditions, the animals were ventilated using a face mask. Following imaging at the awake state, isoflurane was applied at the desired dose (0.5 LORR, 1 LORR or 1.5 LORR). Imaging of the anesthetized state started 12-15 minutes after induction in order to assure arrival to near equilibrium value of the isoflurane (Eger and Johnson, 1987; Wahrenbrock et al., 1974). In addition, at higher LORR (1 or 1.5) we confirmed loss of blinking and withdrawal reflexes, as expected for these states, before imaging started.

### Dataset and conditions

A total of 24 imaging sessions were recorded from 21 mice. Imaging sessions were divided into three groups (1) Awake & 0.5LORR (n=8 mice): these animals were imaged under awake condition and 0.5 LORR i.e. sedation state. (2) Awake & 1 LORR (n=7 mice): these mice were imaged under awake condition and 1 LORR i.e. LOC state. (3) Awake & 1.5 LORR (n=9 mice): these mice were imaged under awake condition and 1.5 LORR i.e. deep anesthesia. For ach awake or LORR condition we collected ~20 trials with visual stimulation and ~20 trials in the blank condition (novisual stimulation; used as control to remove slow trends and the heartbeat artifacts). At the end of anesthesia imaging session, isoflurane was stopped for 15 minutes allowing awakening. Following verification of whisking and reflexes return, a second awake imaging session was performed as a control. In this control we verified that the evoked population responses recovered, and that the change in the VSD signal under anesthesia was not due to dye bleaching or washout over time. In addition, the animals in group (1) were also used to measure the VSD signal throughout the entire range of conscious states in a sequential manner (starting with awake, 0.5 LORR, 1 LORR, 1.5 LORR) within the same animal.

### Visual stimulation

Visual stimuli were displayed on a LCD screen (27×48 cm, 60 Hz refresh rate) positioned at a distance of 15 cm. To stimulate only one eye, the monitor was positioned at ~45^0^ to the long axis of the animal. To measure the visually evoked response we presented to the animal a square stimulus (size of 48 deg) of high-contrast moving gratings (spatial frequency: 0.04 cycles/deg and temporal frequency of 1.5 Hz) over a gray screen. The mean luminance of the grating and gray background was identical. During the inter-trial interval the animal was presented with a gray screen (85 cd/m^2^). The stimulus was position at the center of the screen and presented for 250 ms.

### Optical imaging using voltage-sensitive dyes

We used the MicamUltima system: spatial resolution of 100 × 100 pixels/frame (the whole image covers an area of 5^2^ mm^2^; thus each pixel covers a cortical area of 50^2^ μm^2^) and temporal resolution: of 100 Hz (i.e. frame duration 10 ms). During imaging, the exposed cortex was illuminated using an epi-illumination stage with an appropriate excitation filter (peak transmission 630 nm, width at half height 10 nm) and a dichroic mirror (DRLP 650 nm), both from Omega Optical, Brattleboro, VT, USA. In order to collect the fluorescence and reject stray excitation light, barrier post-filter was placed above the diachronic mirror (RG 665 nm, Schott, Mainz, Germany).

### VSDI analysis

All data analyses were done using MATLAB software. The basic analysis of the VSD signal is detailed elsewhere (Ayzenshtat et al., 2010; Margalit and Slovin, 2018) (see supplementary figure 12 in Ayzenshtat *et al.*, 2010). Briefly, to remove the background fluorescence levels, each pixel was normalized to its baseline fluorescence level (average over first few frames, before stimulation onset). The heart beat artifact and the photo bleaching effect were removed by subtraction of the average of blank signal (recorded in absence of visual stimulation) from stimulated trials. Thus, the imaged signal (Δf/f) reflects the changes in fluorescence relative to the blank trials. For further analysis, VSD signal and maps were computed by averaging over all trials.

### Defining regions of interests (ROIs)

To study the temporal properties of the VSD signal in the different visual cortices, we defined ROIs over the different visual areas. We first computed the averaged activation maps evoked by visual stimulation for each area. For V1 and ESL, maps were averaged at 200-250 ms post stimulus onset and for ESM the mean map average at 300-350 ms post stimulus onset. We then fitted for the awake condition an elliptical shaped ROIs that were fitted to 70% of peak activation pattern for each cortical area, separately. By averaging the VSD signal over pixels within each ROI we obtained the time course (TC) of VSD response for that area.

### Analysis of response latency

To analyze the response latency of the VSD signal in the awake and anesthesia conditions, we selected pixels with significant evoked response to the visual stimulus. A minimal threshold of peak VSD response (averaged across 200-300 ms after stimulus onset) of 3 STDs (relative to baseline activity) was defined. Next, we selected pixels with VSD signal crossing a threshold of 3 STDs (within time window of 50-350 ms from stimulus onset) for at least 4 consecutive time points. We then used linear interpolation on the VSD signal to define the exact time for crossing the threshold. Finally, to compute the latency difference between the awake and different anesthesia conditions we used pixels within the ROI of each area that had valid latency value for both awake and anesthetized condition.

Normalized pixel count difference (NPCD): To compare the number of pixels with valid response latency in each area (Fig 3Ci, Di), for the awake and anesthetized conditions we defined the normalized pixel count difference (NPCD) measure as follows:

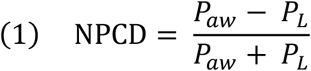

Where P_aw_ and P_L_ are the number of pixels within an ROI that had a valid latency measure in the awake or anesthesia conditions. NPCD=0 means that the number of pixels with valid latency measure was identical on the awake and anesthetized condition. NPCD > 0 means that the number of pixels was higher in the awake condition and NPCD < 0 means higher number of pixels in the anesthetized condition. Normalized latency difference (NLD): To quantify the latency difference between the awake and anesthetize condition (Fig 3Cii, Dii), we defined the measure normalized latency difference (NLD) as follows:

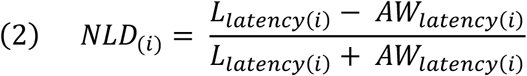

Where L_latency_ and AW_latency_ are the latency of pixel (i) for the anesthesia condition (L; LORR) and awake (AW) condition. Positive values of NLD measure means longer latency values for the anesthetized vs. the awake condition, zero value means no difference between the two conditions and negative values means longer latency values for the awake condition vs. the anesthetized condition. NLD was calculated separately for each pixel that crossed the latency SNR threshold (see latency analysis above). Finally, to assure reasonable SNR, NLD was computed for each area, if the total number of pixels for that area > 150.

### Spatial spread and space-time maps

To quantify the spatial spread of the visually evoked activity and compare the awake and anesthetized conditions, we applied spatial profile on the VSD maps (Fig. 4A inset; width 3-10 pixels) centered on peak response in V1 or ESM or ESM. Next, we averaged the VSD signal across the width of the spatial to obtain the space-time plots of the spatial profile (Fig. 4A) or the spatial profile (i.e. curve; Fig. 4B) at peak response in the awake and anesthetized conditions.

**Figure 1:**
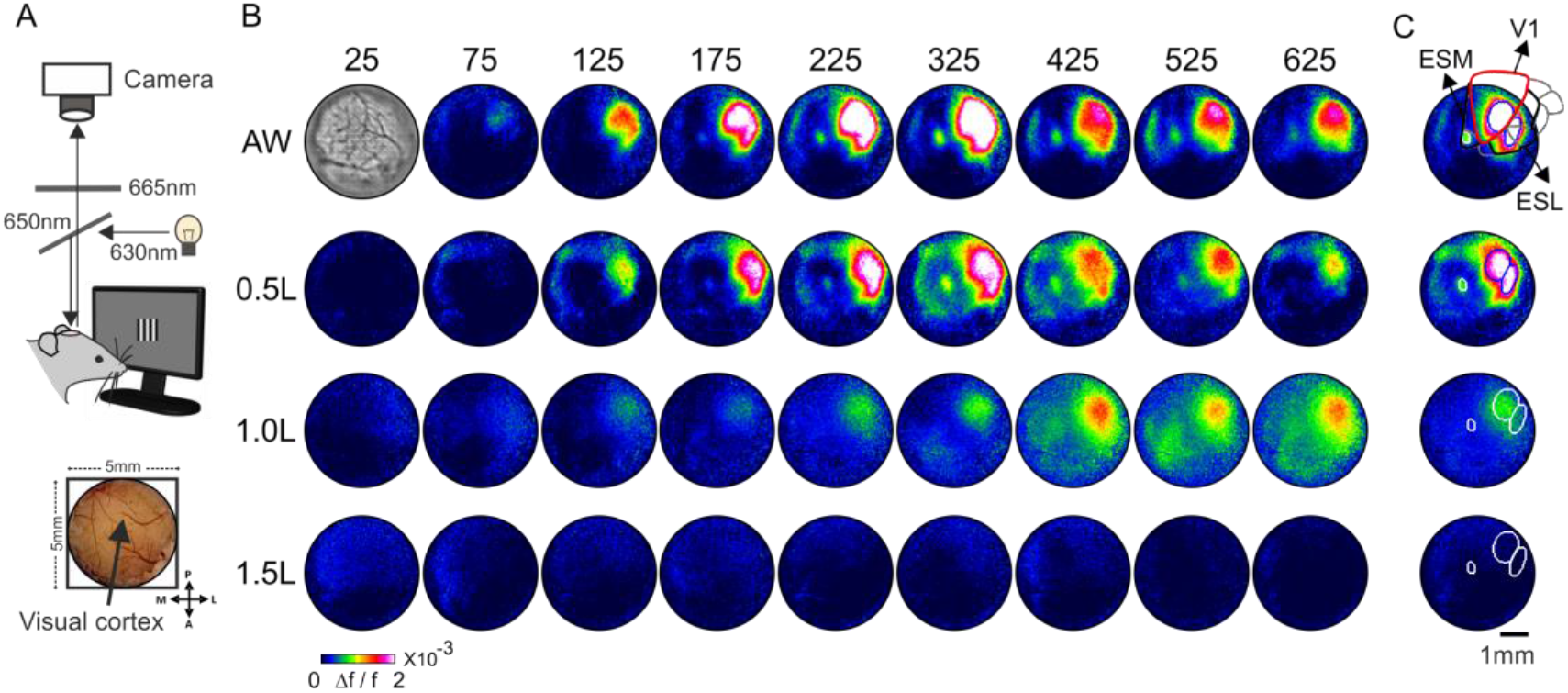
Spatio-temporal patterns of neuronal population in the visual cortex in the awake condition and under increasing isoflurane levels. **A.** Top: A schematic diagram of the VSDI setup combined with visual stimulation. Bottom: An image of blood vessels patterns of the visual cortex. **B.** Population response maps evoked by a square moving grating (see Material and Methods), example session from one animal. The fluorescence level (Δf/f) is depicted by the color bar, each map was averaged over a time window of 50 ms (the numbers above the maps depict the mid time window, after visual stimulus onset). The map at t=25, awake (AW) condition shows the blood vessels pattern of the imaged window. **C:** VSD maps at peak response (averaged over 200-300 ms after stimulus onset) for each condition. The fitted ROIs for each area (see Methods): V1, ESL (extrastriate lateral) and ESM (extrastriate medial) are superimposed on the maps, for each condition: AW and the different anesthesia levels: 0.5 Lorr (0.5L); 1 LORR (1.0L); 1.5 LORR (1.5L). Gray contour on the peak AW map shows the superposition of mouse template for visual cortical areas that was fitted to the imaged window. Red contour denotes area V1 and black contours denotes ESL region and ESM region.

**Figure 2:**
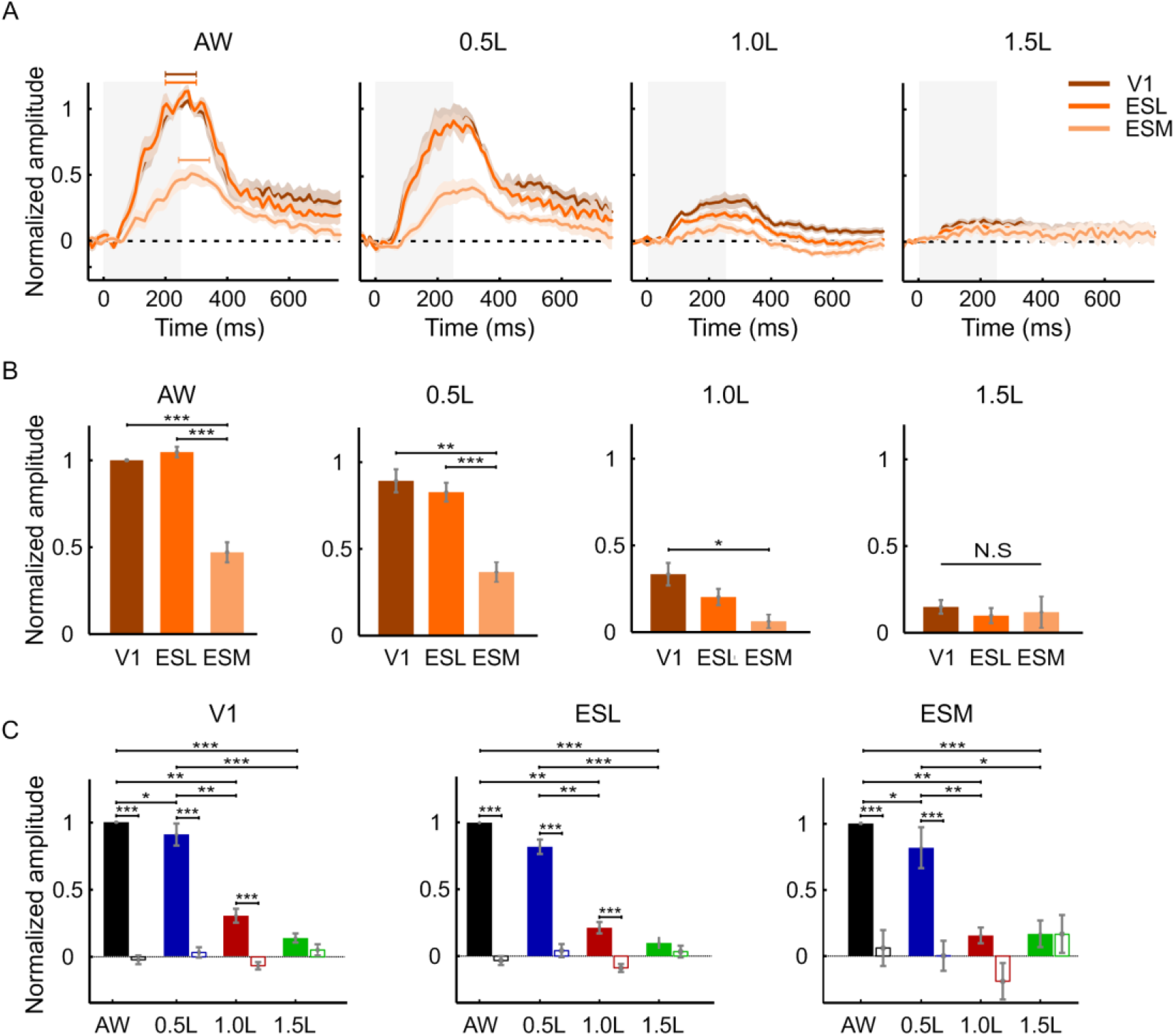
Normalized time course of VSD and peak response evoked by visual stimulation in the awake condition and under increasing isoflurane levels. **A:** Grand average of the normalized time course in the different areas and conditions. In each recording session, the VSD signal was computed for V1 ROI (brown curve), ESL ROI (orange curve) or ESM ROI (light orange curve) and then was normalized to V1 peak response of the awake condition. Next, the time course for each condition was average across all sessions (AW, n=8; 0.5L, n=8; 1.0L, n=7; 1.5L, n=9 sessions). Grey rectangle represent the stimulus duration. Color shaded area around the curves represents ±1 SEM over sessions. **B:** The normalized VSD peak response for the data in (a). Response was averaged over a 100 ms time window (marked with horizontal colored bars, left panel in (a)). Time window for V1 and ESL: 200-300 ms post stimulus onset. ESM: 240-340 ms post stimulus onset. **C:** Peak VSD response normalized to the AW condition of each area. The averaging time window was similar to the time window in (B). Filled bars represent the peak response evoked by visual stimulation and the hollow bars represent the mean VSD response in the blank condition, i.e. no visual stimulation. The response in the blank condition was computed for the same time window as for the visual stimulus condition. Error bars represents ±1 SEM over sessions. Wilcoxon rank-sum test: * p<0.05 ** p<0.01, *** p<0.001. N.S.: non-significant.

**Figure 3:**
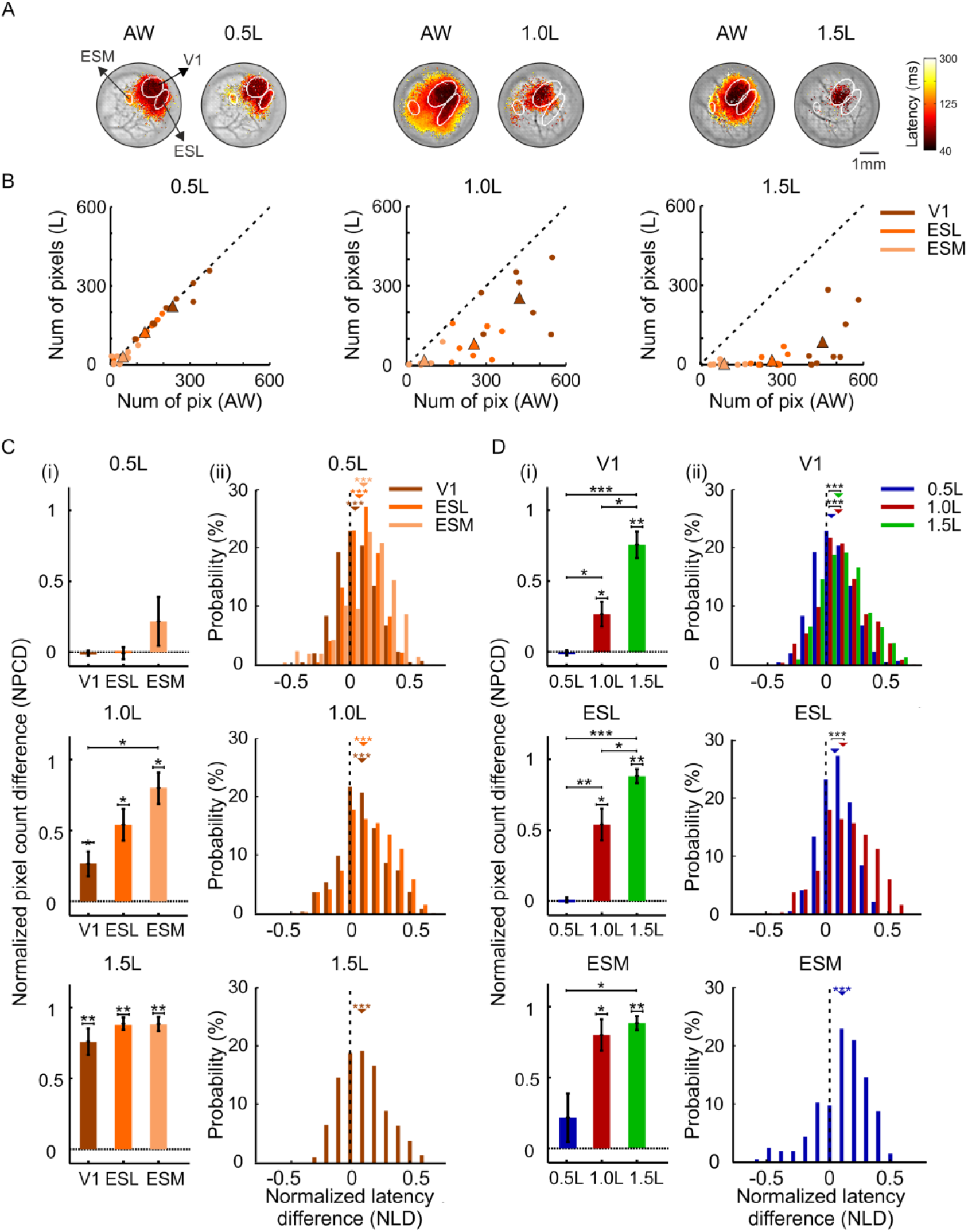
Analysis of the response latency. **A:** Latency maps of visually evoked VSD response in the awake condition and under different isoflurane levels, shown over the blood vessel map of each session. Example sessions from three different animals, white contours represent the ROIs of the different visual areas for each animal. The value of latency onset (see Material and Methods) is color coded as shown by the color bar. **B:** The number of pixels with valid latency value (see Materials and Methods) in the awake vs. anesthetized conditions (left: 0.5L, n = 8; middle: 1.0L, n=7; right: 1.5L, n=9) for each area. Filled circles show the mean (across ROI pixels) value in each recording session. The triangles show the mean value across all recording sessions for each condition. **C:** (i) The normalized pixel count difference (NPCD; see Materials and Methods) under the different isoflurane levels. Top: 0.5L, Middle: 1.0L; Bottom: 1.5L. (ii) Distribution of normalized latency difference (NLD; see Materials and Methods) in the different isoflurane levels: 0.5L (top) 1.0L (middle) and 1.5L (bottom). Small upside-down triangle depicts the mean latency for each condition. **D:** Same data as in (C) but arranged for each area. (i) The NPCD across the different areas and (ii) Distribution of NLD in the different visual areas: V1 (top) ESL (middle) and ESM (bottom).Small upside-down triangles depict the mean latency. Wilcoxon rank-sum/signed-rank tests: * p<0.05, ** p<0.01, *** p<0.001. The number of pixels used for NLD from all sessions: 0.5L: V1 n=1705, ESL n=973, ESM n=205; 1.0L: V1 n=1690; ESL n=554; 1.5L: V1 n=774.

**Figure 4:**
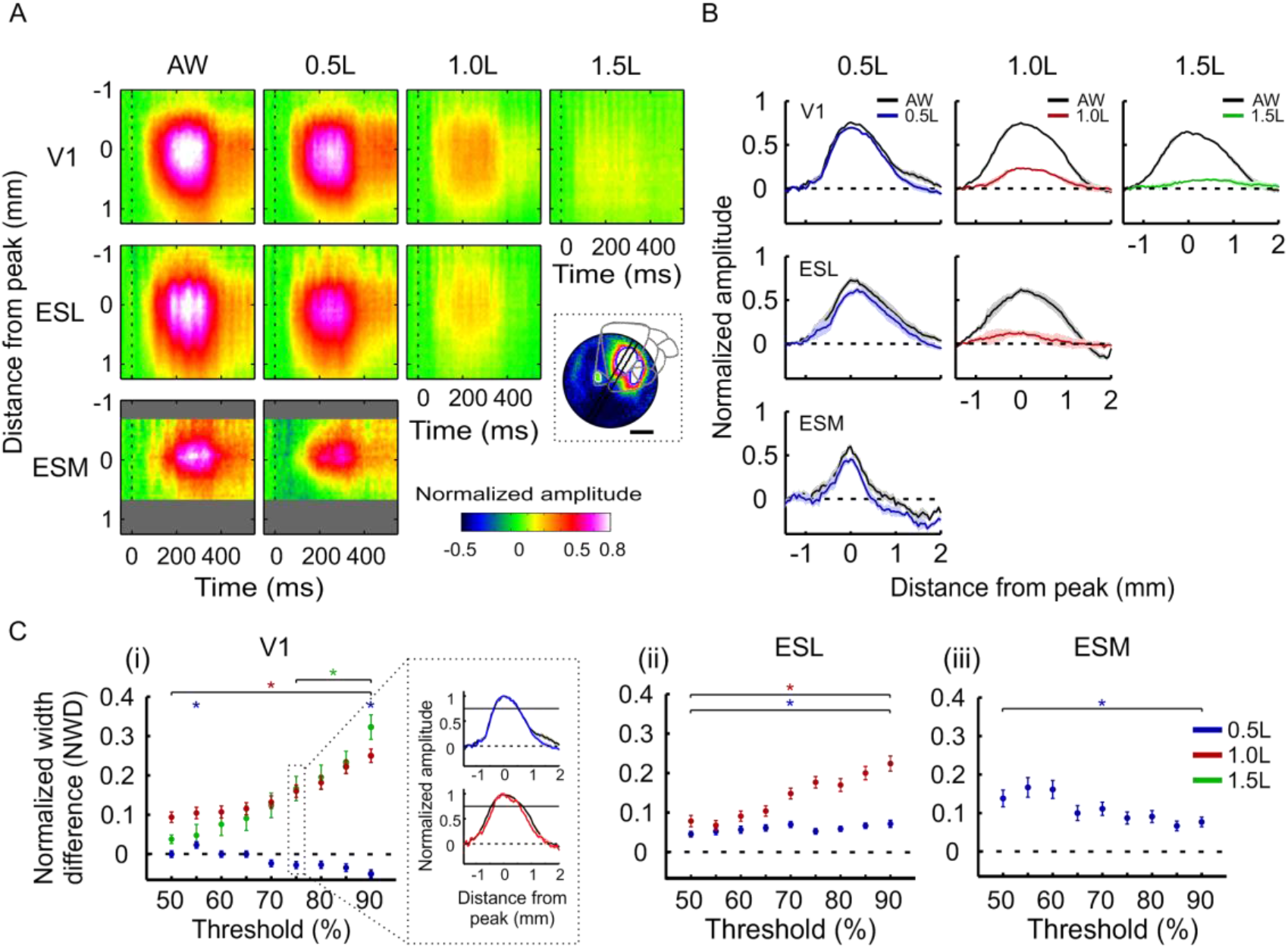
Space-time analysis. **A:** Space vs. time plots: the normalized VSD response at increasing distances from awake peak amplitude (distance=0; y-axis) as function of time (x-axis). The space-time map was computed using a spatial cut that was position over the activation of the analyzed area (inset and see Martials and Methods). The response in each area was normalized to peak activation of the awake condition. **B**: Spatial profile at time of peak response amplitude for each area and condition. The VSD response was normalized to peak response in the awake condition. **C:** Normalized width difference (NWD) in awake vs. anesthetized conditions of increasing thresholds (50 - 90 % from peak response) for V1 (i) ESL (ii) and ESM (iii). Inset in (i) shows the mean spatial profile at 70% from peak response for 0.5L and 1.0L conditions. Wilcoxon signed-rank tests: * p<0.05.

The width of the spatial profile was computed at different percentile (50%-90%) of peak response. For each percentile threshold we found the outer most pixels on each side of the spatial profile crossing the threshold. We then used linear interpolation to find the accurate location of threshold crossing on the spatial profile and also the spatial width. This approach enabled us to estimate the width of the spatial profile at higher accuracy than the used pixel size (50 μm).We then defined a measure for the width difference between the awake and anesthetized condition (Fig. 4C), which we termed as the normalized response width difference (NWD):

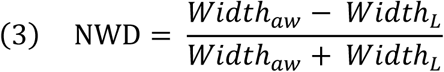

Where Widthaw and WidthL are the width of spatial profile (at various % of peak response, see Fig. 4C) in the awake and anesthesia condition. Positive values of NWD measure means larger width values for the awake vs. the anesthetized condition, zero value means no difference between the two conditions and negative values means larger width values for the anesthetized condition vs. the awake condition. NWD was calculated separately for each session and each time point.

### Correlation analysis and seed correlation maps

To compute synchrony between neural populations, we computed correlation at the single trial level. We first subtracted the mean evoked VSD time course (averaged across trials) from the time course of each pixel (Eq. 4; Brody, 1999). This was done separately for each condition, for awake trials and anesthetized trials. Using this procedure we removed most of the neural activity that is directly related to the visual stimulus onset.

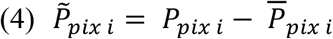

Where P_pix i_ is the population response in pixel i from a single trial and 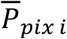 is the mean population response in pixel i across trials (we excluded from the mean response the specific trial). Using a sliding rectangular window of 11 frames (110 ms) we calculated the Pearson correlation coefficient (r; Eq. 5) between each pixel in a seed ROI (3 × 3 pixels, 150 × 150 μm^2^) to all other pixels in the imaged window. The seed ROI was located at the spatial location of peak VSD response for the awake condition. Different seed ROIs were used for V1, ESL and ESM.

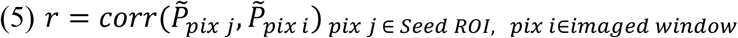

Where corr is Pearson correlation coefficient, 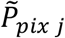 is the VSD signal of pixel j within the seed ROI and 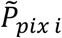 is the VSD response of pixel i in the imaged area. This was computed for each time window width (110 ms) along the time axis: −150 ms to 450 ms from stimulus onset, for each single trial. Next, *Synchrony map* was defined as the mean r values between the seed ROI pixels to all other pixels in the imaged area over a time window (140-300 ms from stimulus onset), for a single trial. Synchrony maps were then averaged across trials, for the same seed ROIs. The outcome of this procedure was a separate synchronization map, for each seed ROI. These correlation maps preserved the anatomical relation between the different areas and revealed the functional correlation between the seed ROI to V1, ESL and ESM (correlation between seed pixels to themselves were excluded from the data).

To evaluate the statistical significance of the seed correlation maps we computed shuffled correlations maps. We calculated the correlation (r) between the seed pixel in a *given trial* to all other pixels in a *randomly chosen trial* of the same condition. This was repeated over 100 permutations to calculate the mean and STD of r values per pixel in the shuffle correlation maps. Next, we set a threshold for correlation (r values were averaged across windows located 140-300 ms post-stimulus onset) as mean±2STDs. Thus, in the real data, only pixels with correlation values exceeding this threshold were further analyzed.

Finally, we calculated the Δ Synchrony between the awake and anesthetized conditions (Fig. 5C) by subtracting the mean synchrony in the anesthetized condition from the mean synchrony in the awake condition (Eq. 6).

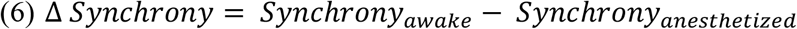

**Figure 5:**
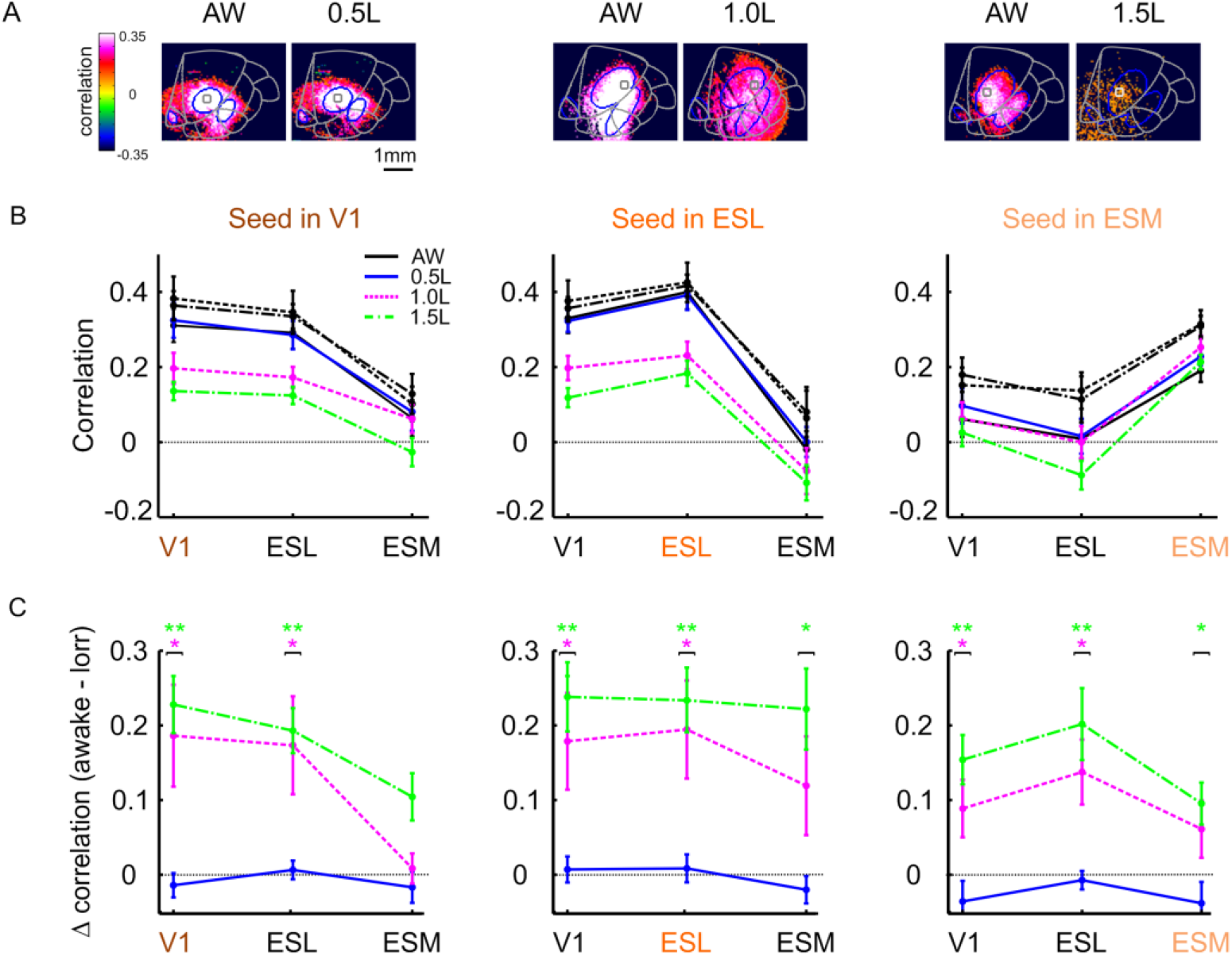
Seed correlation maps in the awake condition and under different isoflurane levels. **A:** Correlation maps, where the seed ROI is located in V1 (small gray squares), in the awake condition (left panels in each pair of maps) and for the different anesthesia levels (right panels in each pair of maps). Three different example sessions are shown from three different animals. Blue contours represent the ROIs of the different visual areas for each animal. Gray contour at AW condition map shows a super position of the mouse template for visual areas fitted to the imaged areas. **B:** Grand analysis of correlation (see Materials and Methods) for the different areas and conditions. For each seed ROI location (V1 – left; ESL – middle; and ESM - right), the correlation values were averaged over a time window (50-400 ms post stimulus onset), and across the pixels of each area and finally across sessions. **C:** Correlation difference (Δ correlation; awake minus anesthetized conditions) over the same time window as in B. Wilcoxon signed-rank tests: * p<0.05, ** p<0.01.

This was done separately for the correlation maps computed for each of the seed ROIs. Positive values of Δ Synchrony means higher correlation for the awake vs. the anesthetized condition, zero value means no difference between the two conditions and negative values means higher correlation for the anesthesia condition. Synchrony was calculated separately for each sessions ROIs (V1, ESL, ESM) and seed pixel area.

### Statistical analysis

To compare the VSD response across different anesthesia states, we used Wilcoxon Rank-Sum and Wilcoxon Signed Rank tests. Bonferonni correction was applied for multiple comparisons. Data are presented as mean ± SEM or median ± MAD.

## Results

We investigated neuronal population responses in mice evoked by visual stimulation under different consciousness levels. Using isoflurane, we controlled the consciousness levels all the way from wakefulness to a deeply anesthetized condition while we imaged the neuronal responses in the visual cortices. We first determined the concentration of isoflurane required to obtain a loss of righting reflex (LORR; see Materials and Methods) for each animal. We then measured the neural population activity from the visual cortex, in four different conditions: wakefulness, mild sedation (0.5 LORR), loss of consciousness (LOC, 1 LORR) and deep anesthesia (1.5 LORR). We used voltage-sensitive dye imaging (VSDI) to measure neural population activity at high spatial (mesoscale, 50^2^ μm^2^/pixel) and temporal resolution (100 Hz; see Materials and Methods) simultaneously. In our experimental setup, the fluorescence dye signal from each pixel reflects the sum of membrane potential from all neuronal elements (dendrites, axons and somata) and therefore is a population signal (rather than response of single neurons). Finally, the VSD signal we measured, emphasizes subthreshold membrane potentials, but reflects also suprathreshold membrane potentials (i.e. spiking activity; see Ayzenshtat et al., 2010; Grinvald and Hildesheim, 2004; Jancke et al., 2004).

### The effect of anesthesia on the amplitude of the visually evoked activity across cortical areas

Figure 1 shows an example session where the VSD signal was measured from the visual cortex of one animal, while it was maintained under different consciousness states. We measured the population response in V1, ESL (comprised of LM, RL and AL areas; LM is considered to be a homologous region to primates V2; Glickfeld et al., 2014) and ESM (comprised of AM and PM areas, that could not be discriminated here; Polack and Contreras, 2012), while the animal was presented with a square stimulus (48 deg) of moving gratings at high contrast (Fig. 1A). Figure 1B shows a sequence of VSD maps for each condition: awake (top row), sedation (0.5 LORR; 2^nd^ row from top), LOC (1 LORR; 3^rd^ row from top) and deep anesthesia (1.5 LORR; bottom row). In the awake condition, shortly after the onset of the visual stimulus, the VSD fluorescence signal (Δf/f) shows increased response, corresponding to depolarization of the population signal, in the caudal part of the imaging chamber (Fig. 1B top row). Two patches of activation appear ~ 100 ms after stimulus onset, one that is located more medially and is corresponding to V1 area and another one that appears more laterally, located with the ESL region (Van Den Bergh et al., 2010; Kalatsky and Stryker, 2003; Rosa and Krubitzer, 1999). In subsequent time frames, the population response amplitude increased and spread horizontally within these two regions, until it seems to merge at the border between the two areas (the vertical meridian). At later times ~170 ms after stimulus onset, a small patch of activation emerged, positioned anterior-medial relative to V1, corresponding to AM area within the ESM region. When the visual stimulus is turned off, the population signal decays slowly towards baseline activity. The VSD map averaged across peak response (200-300 ms after stimulus onset; Fig. 1C, top map) shows clear patches of activation in all three regions, as denoted by the visual areal template (adapted from Marshel et al., 2011; V1 is depicted by the red contour; ESM and ESL by the black contour). A similar activation pattern but with a lower response amplitude, appears under 0.5 LORR (Fig. 1B 2^nd^ row from top), while in this condition we can still observe visually evoked response in all three areas (V1, ESL and ESM). Similarly, the VSD map averaged at peak response, shows activation patches in all three regions (Fig. 1C, 2^nd^ map from top). This spatial patterns of activation is dramatically modified in LOC, i.e. 1 LORR (Fig. 1B 3^rd^ row from top), where ESM area shows no evoked response and we can observe a large decrease of the evoked activity in V1 and ESL. This is also evident in the VSD map averaged at peak response (Fig. 1C 3^rd^ map from top) under 1 LORR. Under deep anesthesia (1.5 LORR condition; Fig 1B bottom row) the VSD response maps show no clear patches of activations (Fig. 1C bottom). The results in the example session suggest that the visually evoked responses show a dramatic decrease under 1LORR, i.e. the LOC state.

Figure 2 shows the grand analysis (averaged across all sessions) of the VSD time course (TC) in the different visual areas. Figure 2A depicts the TC (normalized to V1 peak response of the awake condition; see Materials and Methods) for the three cortical regions and for the different consciousness levels. Figure 2A left shows the visually evoked population response in the awake condition: whereas V1 and ESL have similar dynamics and response amplitude, the population response in area ESM seems to lag behind, revealing slower dynamics and a much lower mean peak response amplitude of 47.2% (±5.7, sem) relative to V1; p < 0.001; Wilcoxon rank sum test, Fig. 2B left). The visually evoked responses under 0.5 LORR (sedated state) show a small decrease in peak response amplitude relative to the awake condition (Fig. 2A-B) while the dynamics of the evoked response were generally preserved in the different regions. Figure 2C shows the VSD signal at peak response for each area, when the response of each area was normalized to its own awake condition. In the sedated state (0.5 LORR), the mean V1 peak response decreased to 91% (± 8.24, sem; p<0.05, Wilcoxon rank sum test) of the awake response, the mean ESL peak response decreased to 82% (± 5.4, sem; n.s. (p=0.052), Wilcoxon rank sum test) of the awake response and the mean ESM peak response decreased to 81.7% (±15.4, sem; p<0.05, Wilcoxon rank sum test) of the awake response. Thus, under 0.5 LORR the response amplitude decreased moderately and this not statistically significant in all areas.

A dramatic decrease occurred at LOC, under 1 LORR, which is in accordance with the example session shown in Fig. 1B-C. The grand analysis in Figure 2A-B shows that the VSD response in all areas dropped by 70-85% relative to the awake condition. The mean population response in V1 dropped to 30.14% (±5.2, sem; p < 0.01, Wilcoxon rank sum test; Fig. 2C left) relative to V1 awake condition, the response in ESL dropped to 21.2% (±4.3, sem; p < 0.01; Wilcoxon rank sum test; Fig. 2C middle) relative to ESL awake condition and the response in area ESM decreased to 15.5% (±5.9, sem; p < 0.05; Wilcoxon rank sum test; Fig. 2C right) relative to ESM awake condition. While in the LOC state all areas showed a dramatic reduction of the visually evoked response amplitude, the population responses in V1 and ESL were significantly higher than the no visual stimulation condition i.e. blank condition (all statistical p values (ps) < 0.05 for V1 and ESL; Wilcoxon rank sum test). This means that under 1 LORR, V1 and ESL preserved visual processing capabilities. In contrast, the visually evoked response amplitude in area ESM was not significantly different from blank condition (i.e. no visual stimulation; p=0.065) under 1LORR condition. Thus LOC is characterized by a dramatic decrease of visually evoked responses in all areas, and specifically in ESM, which is considered to be a higher visual area compared with V1 and ESL (both V1 and ESL project to ESM; Glickfeld et al., 2014; Katzner and Weigelt, 2013). This may suggest that higher areas have a higher sensitivity to anesthesia levels, potentially because their visually evoked response amplitude are smaller.

At 1.5 LORR condition (deep anesthesia), all areas show additional decrease in the visually evoked response (Fig. 2A-B right panels). The mean population response in V1 dropped to 13.4% (± 3.4, sem in V1, p < 0.001; Wilcoxon rank sum test; Fig. 2C left panel) of the V1 awake response, the mean VSD response in area ESL dropped to 9.8% (± 3.64 sem; p < 0.001; Wilcoxon rank sum test; Fig. 2C middle panel) of the ESL awake response and the mean VSD response in area ESM was 16.7% (±10.05 p < 0.001; Wilcoxon rank sum test; Fig. 2C right panel) of the ESM awake response. Importantly, the population response under 1.5 LORR was not statistically different from the blank condition for all areas (all ps > 0.1) and in addition there was no statistical difference between the peak response amplitude across the different areas (p > 0.5; Wilcoxon rank sum test). This means that under 1.5 LORR, when averaging the population responses over the entire area of each region: V1, ESL and ESM, the visually evoked responses were at noise level. However, additional analyses we performed below – showed weak neural responses under 1.5 LOOR in some of V1 pixels.

### The effect of anesthesia on the latency of the visually evoked activity across cortical areas

Previous studies that investigated the latency to response onset reported mixed effects of anesthesia: while many studies reported increased latencies, other reported no effect (Sellers et al., 2015) or even decreased latency (Durand et al., 2016). Thus we decided to investigate the response latency of the visually evoked activation and compare it over consciousness levels. Figure 3A depicts latency maps from three example sessions (taken from three different animals), where each pair of maps (awake and LORR condition) shows the response latency value for each pixel within the imaged areas: V1, ESL and ESM. Latencies were computed only for pixels with an SNR threshold (see Materials and Methods) which we termed pixel with valid latency value. Figure 3A left shows the latency maps in the awake and 0.5 LORR (sedated) conditions: there is a small decrease in the number of pixels (with valid latency value) under 0.5 LORR and a minor increase in latency values (latencies are color coded). A dramatic change occurs when switching to 1 LORR (LOC, Fig. 3A, middle): the number of pixels (with valid latency) is dramatically reduced and the latencies values are longer. These effects are even more pronounced under 1.5 LORR (deep anesthesia) as can be seen in Figure 3A right: only a very small number of pixels crossed the threshold of valid latency and the latencies clearly show longer values. Figure 3B shows the grand analysis (across all recording sessions) for the number of pixels at different isoflurane levels. The scatter plots in Figure 3B depict the numbers of pixels under isoflurane (y-axis; 0.5, 1.0 or 1.5LORR) vs. the awake condition (x-axis) for each imaging session (small circles). This is plotted for the different areas (color coded). Under 0.5 LORR, it can be seen that most of the data points are located closely along the diagonal, indicating almost no change relative to awake condition (Fig. 3B left panel). To quantify the change in number of pixels, we calculated the normalized pixel count difference (NPCD, see Material and Methods, Eq. 1) shown on Figure 3Ci for the different isoflurane levels. NPCD=0 means that the number of pixels with valid latency measure was identical on the awake and anesthetized condition. NPCD > 0 means that the number of pixels was higher in the awake condition and NPCD < 0 means higher number of pixels in the anesthetized condition. Figure 3Ci top, shows the NPCD under 0.5 LORR with a non-significant change relative to the awake condition, for all areas (all ps > 0.1; Wilcoxon signed-ranked test). The middle panel in Figure 3B shows the scatter plot for the 1 LORR with a large and significant decrease in the number of pixels in all areas (colored triangles shows the mean pixel count for each area). This is further quantified by the NPCD measure in Figure 3Ci middle panel, which shows positive values, meaning larger number of pixels in the awake condition, in all areas. The mean NPCD in V1 was 0.26(±0.09 sem), in ESL 0.55(±0.11 sem) and in ESM 0.83(±0.11 sem), all ps < 0.05 (Wilcxon signed-rank test). Figure 3B right shows the scatter plot for the 1.5 LORR with an additional decrease in the number of pixels in all areas which is quantified by NPCD in Figure 3Ci bottom panel. The mean NPCD in V1 was 0.75(±0.094, sem), in ESL 0.89(±0.04, sem) and in ESM 0.92(±0.05, sem), all ps < 0.01 (Wilcxon signed-rank test). In summary, this analysis shows a dramatic decrease in the number of activated pixels within each area, under LOC and then deep anesthesia. Only a small percentage of pixels (i.e. small part of neuronal population) show valid latency responses in V1, ESL and ESM in the latter condition.

Next, we quantified the changes in latency values under the different isoflurane levels, for each area separately. We computed the normalized latency difference (NLD; see Materials and Methods, Eq. 2) separately for each pixel (with valid latency value) under each LORR relative to the awake condition. NLD=0 means no latency difference between the awake and anesthetized states. NLD > 0 means that the latencies were longer in the anesthetized state and NLD < 0 means longer latencies values in the awake condition. The distribution of NLD values in Figure 3Cii depicts the latency difference in all pixels for the different areas (color coded). Already under 0.5 LORR (Fig. 3Cii top) there is a clear shift of the NLD distribution towards positive values, with higher areas showing larger mean NLDs. The mean NLD value under 0.5 LORR for V1 is 0.04(±0.004, sem), for ESL: 0.07(±0.005, sem) and for ESM: 0.1(±0.007, sem), all ps < 0.001 (Wilcxon signed-rank test). Under 1 LORR (Fig. 3Cii middle), there is a further increase in the NLD with a mean value in V1 of 0.1(±0.005, sem) and for ESL: 0.15(±0.01, sem) all ps < 0.001 (Wilcxon signed-rank test; we note that the NLD in area ESM under 1 and 1.5 LORR could not be computed due to a low number of pixels as well as for area ESL under 1.5 LORR). Finally under 1.5 LORR (Fig. 3Cii bottom), V1 showed an NLD value of 0.1(±0.007, sem), p < 0.001 (Wilcxon signed-rank test).

Figure 3D shows the NPCD values and NLD distributions within each area, under different isoflurane levels. While area V1 shows a gradual increase in the NPCD with increasing levels of isoflurane (Fig. 3Di, top), area ESM shows a large NPCD value already at 1 LORR (Fig. 3Di bottom). The NPCD in area ESL shows in-between values (Fig. 3Di middle). Figure 3Dii shows the distribution of latencies values as quantified by NLD, for each area, allowing a clear view of how the increased isoflurane level affects the NLD values within each area, shifting it towards more positive values under higher levels of isoflurane. In area V1 the mean NLD values increased by a factor of 2.4 from 0.5 to 1.5 LORR (p<0.001, Wilcoxon rank sum test). Area ESL showed a similar phenomenon and the mean NLD significantly increased from 0.5 to 1 LORR (p=0.002) by a factor of 1.95. Finally, area ESM showed the largest NLD value in 0.5 LORR, the mean value was 0.1(±0.01, sem; p<0.001 Wilcoxon signed-rank test). In summary, the above results suggest that area ESM shows the highest sensitivity to isoflurane levels: larger values for NPCD (p < 0.05 for V1 and ESM comparison under LOC) and NLD at lower isoflurane levels relative to areas V1 and ESL (p <0.001 for ESM comparison with V1 and ESL under sedation).

### The influence of anesthesia on the local spatial spread of visually evoked response

The horizontal spread of local visual stimuli within V1 was suggested to be mediated by horizontal connections located mainly in the upper layers (Fehérvári et al., 2015; Stettler et al., 2002). Thus, our next step was to study how the spatial spread of the visually evoked activity, is influenced by isoflurane levels. To study this, we computed a spatial cut (Fig. 4A, inset, black spatial cut; see Materials and Methods) crossing trough peak response activation in V1, and additional separate spatial profiles crossing through peak response in ESM and ESL. Computing the spatial profile (by averaging over the width of the spatial cut) as function of time results in space-time maps which are depicted separately for V1, ESL and ESM. Figure 4A shows space-time maps the grand analysis of i.e. the mean across all recording sessions, for awake and each of the isoflurane levels. The space-time maps (zero on the y-axis depicts the peak spatial response in each region) clearly show that the evoked response in the awake condition spreads laterally in V1 and ESL, while in area ESM the spatial spreads is much smaller. Under increasing levels of isoflurane, the spread of the visually evoked response is reduced in all regions. To quantify this, we computed the mean (across sessions) spatial profile curves, aligned on the spatial location of peak response, for each area and condition (Fig. 4B). Interestingly, under sedation (0.5 LORR), area V1 did not show a clear change in the horizontal spread, however the width of the spatial profiles of both ESL and ESM decreased and this was accompanied by a small reduction of the response amplitude (Fig. 4B, left column). Under LOC (1 LORR), the peak response in V1 and ESL decreased dramatically while accompanied by a decreased spread of activation (Fig. 4A-B, middle column; the response amplitude of area ESM under 1 LORR, was n.s. from the blank condition and thus the space-time map is not shown). Under 1.5 LORR, area V1 shows a very weak response and a much smaller spread of activation. To investigate whether the decrease in the spatial spread resulted not only from decreased peak response, but also from a change in the shape of the spatial profile we computed the normalized width difference (NWD) which is a measure for the width of the spatial spread (see Materials and Methods Eq. 3). NWD = 0 means no difference between the awake and anesthetized conditions, NWD > 0 means larger width values for the awake vs. the anesthetized condition, and NWD < 0 means larger width values for the anesthetized vs. the awake states. NWD was calculated for percentile values ranging from 50%-90% of peak response for each condition and area. Figure 4C shows that while under 0.5 LORR V1 shows mostly no significant change in NWD, areas ESL and ESM already show a significant decrease for most peak percentiles (Fig. 4Ci blue dots vs. Fig. 4Cii and iii blue dots). The median NWD in V1 for 0.5 LORR varied from −0.05(±0.01 mad) to 0.02(±0.008 mad) and was not statistically significant for most percentile points. The median NWD for ESL in 0.5 LORR varied from 0.045(±0.007 mad) to 0.07(±0.007 mad) p < 0.05 (Wilcoxon signed-rank test) for all percentile points. This is also the case for area ESM, where the median NWD varied from 0.06(±0.012 mad) to 0.17(±0.026 mad), p for all percentiles < 0.05 (Wilcoxon signed-rank test).

Interestingly, under 1 LORR condition, both V1 and ESL showed a larger change in the median value of NWD which varied in V1 from 0.09(±0.01 mad) to 0.25(±0.017 mad) and in ESL: from 0.07(±0.012 mad) to 0.22(±0.018 mad). Finally, under 1.5 LORR NWD could be computed only for area V1 and the median values varied from 0.04(±0.01 mad) to 0.32(±0.03 mad); p < 0.05 for percentiles > =70. In summary, the results show that the horizontal spread of visually evoked response in V1 is smaller, in a dose response manner which further suggests that isoflurane impairs the information propagation within horizontal connections in V1. A similar conclusion can be drawn for area ESM which in our set of experiments, is comprised mainly of area AM. The spatial spread was narrower under increasing levels of isoflurane also in ESL, however, this region is likely to reflect the activity from few separate visual areas and thus the spatial spread reduction can reflect information flow between the these areas. Finally, area ESM showed the largest NWD values, already under 0.5 LORR (p < 0.01 for V1 and ESM comparison), suggesting larger sensitivity of ESM to isoflurane.

### The effect of anesthesia on intra-areal and inter-areal synchronization

Intra-areal and inter-areal synchronization is an important neural mechanisms of visual processing. To study how anesthesia influences cortical synchronization involved in visual processing, we computed synchronization maps at the single trial level (see Materials and Methods Eq. 4–6). Using a sliding window of 110 ms we calculated the Pearson correlation coefficient (r) for all pixel pairs *between* the seed ROIs and all other pixels within the imaged visual cortex. This was done for each trial separately, after removing the direct stimulus contribution by subtracting the mean (across trials) visually evoked response from each pixel (Eq. 4 in the Materials and Methods). Next, the correlation maps were averaged across trials and for all seed pixels, which yielded spatial synchronization map as depicted in Figure 5A. The synchronization maps show the zero-time lag correlation between the seed pixels and the pixels in all other areas, during visual stimulus presentation (Gilad and Slovin, 2015; Maatuf et al., 2016). We further analyzed only pixels with correlation values exceeding the mean shuffle condition by 2STDs (see Materials and Methods). Thus, the resulted correlation maps show the population synchronization (functional connectivity; as the contribution of the visual stimulus was subtracted) at the pixel level, between the seed ROI to all other imaged pixels. Importantly, this analysis enables not only to study the functional strength of the correlations (r) but also their anatomical relation.

Figure 5A shows examples of correlation maps in three different animals, where the seed ROI is located in V1 (small gray squares; seed ROI is positioned over the spatial peak response in V1). For each animal the awake condition is shown along with the relevant anesthesia condition. Figure 5A left shows the correlation maps in the awake and 0.5 LORR condition of the same animal. Clear patches of higher correlation (white pixels) appear in V1, ESL and ESM under the awake condition. Interestingly, these patches of functional correlation are highly similar to the patterns generated by axonal projections stained by local marker injections in V1 to ESL and ESM (Wang et al., 2012). A similar correlation map emerged under 0.5 LORR – which shows little change between the awake and sedated state. Figure 5A middle panels shows the correlation maps in awake and under 1 LORR. Again, under LOC there is a large change of the neuronal measure i.e. decrease in correlation. Figure 5A right shows the correlation maps in the awake and under deep anesthesia: the correlation values further decrease to almost zero.

Figure 5B shows the grand analysis across all recording sessions and plots the mean correlation values in the awake condition (black curves) and under the different isoflurane levels (colored curves). The curves show the correlation values in the different areas, when the seed ROI is in V1 (Fig 5B, left), in ESL (Fig 5B, middle) and in ESM (Fig. 5B, right). Interestingly, in the awake condition, the intra-areal correlation shows the highest values, i.e. the correlation values between the seed and all other pixels within the same area. The mean intra-areal correlation values in the awake condition in V1 and ESL are comparable (~ 0.35-0.43; difference is n.s.) while the mean intra-areal correlation in area ESM is 0.28 which is significantly lower than that in V1 and ESL (all ps < 0.05; Wilcoxon rank sum test). The inter-areal correlations (e.g. between seed ROI in V1 to ESL/ESM; between seed ROI in ESL to V1 and ESM etc) in the awake condition shows typically lower correlation compared with intra-area correlation. The synchronization between V1 and ESL in the awake condition has high values, while the functional connectivity between V1 to ESM is lower, which is again in accordance with known anatomical connectivity (Wang et al., 2012).

Figure 5B shows that under 0.5 LORR (blue curves) there is no significant change of correlation relative to the awake condition, either locally, i.e. within an area or between areas. This is further quantified by the delta correlation, awake minus anesthesia condition, as shown in Fig. 5C (blue curves; n.s. from zero). Interestingly, under 1 LORR (pink curves, Fig. 5B-C), we see a dramatic decrease in correlation values for most cases, a phenomenon that is even more pronounced under 1.5 LORR (green curves; Fig. 5B-C). Both V1 and ESL show a decrease of ~50% in intra-areal correlation under 1 and 1.5 LORR. Under 1 and 1.5 LORR, the inter-areal synchronization between seed ROI in V1 and ESL is also reduced by ~ 50% as well as the correlation between the seed pixels in ESL and V1. Interestingly, the inter-areal correlation between area ESM and V1 or ESL, under 1 and 1.5 LORR drops to zero (values that are non-significant from zero). These results suggest that are ESM becomes disconnected from the visual network under LOC and also deep anesthesia. In summary, isoflurane affects both intra-areal and inter-areal synchronization. At higher doses of isoflurane, ESM seems to be completely “disconnected” from the network, while the correlation between V1 and ESL are dramatically reduced, but still remains larger than zero. Finally, high doses of isoflurane partially spare intra-areal synchronization while affecting more dramatically inter-cortical connections.

## Discussion

We characterized the spatio-temporal patterns of neuronal responses evoked by visual stimulation under varying concentrations of isoflurane, from wakefulness to deep anesthesia. Our results extend previous electrophysiological studies, by measuring population responses at high spatial (meso-scale) and temporal resolution from several visual areas simultaneously. We found that isoflurane has multiple effects on visual processing: from population responses to network synchronization and these effects are dose dependent.

### Reduced neural response and slower dynamics under higher isoflurane levels

We found that the response amplitude evoked by visual stimulus decreased under isoflurane. This effect was larger at higher isoflurane doses. This replicates previous studies reporting of an overall reduction in neural activity during anesthesia (Aasebø et al., 2017; Hentschke et al., 2005; Vaiceliunaite et al., 2013) and it is also in agreement with previous VSDI studies in the barrel cortex of awake versus anesthetized mouse (Devonshire et al., 2010; Ferezou et al., 2006). Depending on the anesthetic agent, dose and combination with additional drugs (sedative or analgesic) and the response to sensory stimulation decreased by 30-60% under anesthesia. In our study, the response amplitude was relatively preserved under sedative concentrations of isoflurane, i.e. the response decreased by 9-19 % in the different regions. This is in agreement with studies using low to intermediate isoflurane levels (Berger et al., 2007; Sellers et al., 2015) that reported preserved or minimally decreased neural responses. A reduction of 70-85% of the response amplitude was observed at higher isoflurane levels inducing LOC (Aasebø et al., 2017).

Increased latencies of visually evoked responser were found in V1 under different anesthesia protocols, including isoflurane and for different visual stimuli (Aasebø et al., 2017; Durand et al., 2016; Gao et al., 2010; Marshel et al., 2011; Pisauro et al., 2013). This is in accordance with our results of increased population response latency in all areas under anesthesia. In contrast, Sellers et al. (2015) reported no alteration in V1 response latency under increasing isoflurane dose (0.5%-1%), however, this study combined xylazine with isoflurane in ferrets.

### The spatial spread of the VSD signal in V1 and higher visual areas

We found that the spatial spread of visually evoked population responses in V1 decreased under isoflurane anesthesia dose dependently (Fig. 4). As noted in the Introduction, isoflurane augments inhibitory synapses and depresses excitatory synapses, hence damping network excitability (Campagna et al., 2003; Liang et al., 2012a; Mashour, 2005; Nallasamy and Tsao, 2011). This mechanism can explain the reduced spatial spread reported in numerous studies some of which used isoflurane (Ferezou et al., 2006; Hentschke et al., 2017; Liang et al., 2012b; Mashour, 2005; Raz et al., 2014), including ours. It was recently shown that isoflurane suppresses topdown excitatory pathways, which can further reduce neural population excitability (Hentschke et al., 2017; Liu et al., 2011). The narrowing of spatial spread was more pronounced in ESL and ESM than V1. However, while the ESM region in our study was comprised mainly from area AM, the ESL region is likely comprised from several different visual areas. Thus, while the spatial profile in V1 reflects the propagation via horizontal connections, the ESL results may include propagation between different visual areas as well.

### Reduced local and inter-areal synchronization under different levels of isoflurane anesthesia

Information flow and synchronization within and between brain areas is an important mechanism of sensory processing. This process was also linked to LOC during anesthesia (Aru et al., 2019; Hudetz, 2012; Pullon et al., 2020). By computing synchronization maps when the seed pixels are located in V1 or in higher visual areas (ESL and ESM) we investigated intra-inter-areal functional correlation. Synchronization analysis showed highest values for intra-areal i.e. within V1, ESL or ESM. In addition, feed-forward synchronization was highest between V1 to ESL. Feed-back synchronization from ESL was largest to V1 while ESM originated synchronization was similar to V1 and ESL. Importantly, these correlation strength are in accordance with the anatomical study of Wang et al., 2012 who measured anatomical projections in visual cortices of the mouse. Interestingly, anesthesia reduced the within and between areal synchronization (Fig. 5) but at high isoflurane concentration, ESM showed no correlation with any other areas except itself.

Several previous studies reported decreased brain correlations under anesthesia. A recent study showed that isoflurane (2%) induced decorrelated activity between hippocampal neurons in a mouse (Yang et al., 2020). Furthermore, Liang et al., (2012a) showed that whole-brain functional connectivity strength, measured using BOLD signal in awake and isoflurane anesthetized rats, decreased under anesthesia. Interestingly, they showed that functional connectivity is selectively reduced for regions that are separated by less than 10mm. Unlike the VSD signal that directly reflect neuronal population activity, BOLD signal is an indirect measure of neural response comprised mainly from slow hemodynamic signal which shows a neurovascular decoupling at high isoflurane doses (~1.5%; Masamoto and Kanno, 2012; Schummers et al., 2008; Wu et al., 2016). Another study in ferrets, reported that the amount of local information transfer from V1 to prefrontal cortex is reduced under isoflurane and ketamine administration (Wollstadt et al., 2017). The reduction of local information in V1 might be related to the reduction of within areal correlation we observed. Finally, Pullon et al., (2020) reported that propofol anesthesia induced LOC that was associated with a substantial global decrease in connectivity.

A large number of studies reported increased correlations during anesthesia (Aasebø et al., 2017; Goltstein et al., 2015; Greenberg et al., 2008). We suggest that the inconsistency in correlation results may depend on the anesthesia protocol, experimental protocol (e.g. stimuli) and type of correlation measured. The neural network state (spontaneous vs. evoked response) may later correlation as well. For example, Goltstein et al. (2015) showed that V1 correlations in the spontaneous state are increased under anesthesia, but the overall correlation during visually evoked response did not change. Aasebø et al. (2017) compared single unit correlation in V1 between awake and isoflurane (1%). Unlike our results, they found that visually evoked pair-wise correlations increased under anesthesia. There are several possible reasons for this contradiction. First, we used head fixed mice only, assuring an identical visual stimulation under all conditions, whereas Aasebø et al. used freely moving rats while awake and head fixed under anesthesia. This enforces different visual stimulation during the two states. Furthermore, it was previously shown that locomotion increases visually evoked responses in the visual cortex (Ayaz et al., 2013) and thus can influence the correlation of single units. Second, we subtracted the mean evoked VSD response, which removed most of the activity directly driven by the visual stimulus. Another well studied measure of synchrony is coherence, which reflects synchronization in the frequency domain. Like pair-wise neuronal correlations, coherence analysis is entangled. Some showed enhanced coherence during anesthesia (Michelson and Kozai, 2018; Sellers et al., 2015) while others showed reduced coherence at alpha and frequencies (alpha 8-15 Hz; Redinbaugh et al., 2020) which dominates the VSD signal (Gilad et al., 2012). It has to be noted that coherence studies are strongly affected by the frequency band studied and thus the results may vary depending on the analysis.

### ESM is more sensitive to isoflurane than V1

All neural measures used to quantify the population response: peak amplitude, latency and width of spatial profile, showed the greatest changes in ESM, usually at low isoflurane dose. Under 0.5 LORR, ESM showed the largest latency increase (NLD) and spatial spread decrease (NWD). In addition, area ESM showed no significant evoked response at 1 LORR (while area V1 and ESL still showed a significant response) as well as the largest reduction in the number of pixels with valid latency values (the highest value of NPCD). Interestingly, under higher isoflurane doses, ESM showed decreased correlation with other areas, while V1 and ESL preserved significant synchronization.

There are several possible explanations to the higher sensitivity of area ESM to isoflurane: First, as seen in the awake animal, the population response to the visual stimuli was smaller compared with V1 and ESL (Polack and Contreras, 2012). Hence, a reduction in activity caused by isoflurane at LOC-inducing concentration, will nullify the weaker responses measured in higher visual areas, while only lowering the response in other areas that get direct visual input from the thalamus. The latter is only one synapse away from the retina, and thus remains an effective driving force for V1 activity, even with deeper anesthesia. Second, activity progression within the visual cortex is mediated by cortico-cortical connections (Bienkowski et al., 2019; Fehérvári and Yagi, 2016; Polack and Contreras, 2012; Wang and Burkhalter, 2007; Wang et al., 2011) which were shown to be interrupted with increasing isoflurane dose (Alkire et al., 2008; Raz et al., 2014; see also Discussion on synchronization). Third, the VSDI signal emphasizes subthreshold activity. In lower cortical visual areas this reflects subcortical inputs. A reduction in cortical APs with increasing isoflurane dose (Hentschke et al., 2017), may be less noticeable in VSD signal of lower order areas, but diminish the signal in higher order areas. Thus, we suggest that as isoflurane concentrations increases, higher visual areas are turned off first, whereas lower visual areas become inactive only at higher doses. A recent study by Krom et al. has shown similar results in the auditory system (Krom et al., 2020).

Finally, a number of theories explain consciousness and how anesthesia modulates it. The global neuronal workspace theory (GNW, Mashour et al., 2020) and higher order theories (HOT, Brown et al., 2019; Lau and Rosenthal, 2011) both emphasis the role of higher order cortices (mostly frontal cortex) in creating consciousness, either through making the information globally available (GNW) or by creating the perception required to assign meaning to sensory information. These theories predict anesthetics will preferentially affect higher order areas or top-down pathways arising from them. Other theories, such as information integration theory (Oizumi et al., 2014) and recurrent processing theory (Lamme, 2006), assign a similar weight to top-down and bottom-up pathways. Thus, these theories do not predict a stronger effect of anesthetics on higher order areas. Since we found a stronger effect of anesthesia on higher order areas, our results tend to support the theories requiring this (i.e. GNW & HOT).

## References

Aasebø, I.E.J., Lepperød, M.E., Stavrinou, M., Nøkkevangen, S., Einevoll, G., Hafting, T., and Fyhn, M. (2017). Temporal Processing in the Visual Cortex of the Awake and Anesthetized Rat. Eneuro 4, ENEURO.0059–17.2017.

Alkire, M.T., Haier, R.J., and Fallon, J.H. (2000). Toward a Unified Theory of Narcosis: Brain Imaging Evidence for a Thalamocortical Switch as the Neurophysiologic Basis of Anesthetic-Induced Unconsciousness. Conscious. Cogn. 9, 370–386.

Alkire, M.T., Hudetz, A.G., and Tononi, G. (2008). Consciousness and Anesthesia. Science 322, 876–880.

Andermann, M.L., Kerlin, A.M., Roumis, D.K., Glickfeld, L.L., and Reid, R.C. (2011). Functional specialization of mouse higher visual cortical areas. Neuron 72, 1025–1039.

Aru, J., Suzuki, M., Rutiku, R., Larkum, M.E., and Bachmann, T. (2019). Coupling the State and Contents of Consciousness. Front. Syst. Neurosci. 13, 1–9.

Ayaz, A., Saleem, A.B., Schölvinck, M.L., and Carandini, M. (2013). Locomotion controls spatial integration in mouse visual cortex. Curr. Biol. 23, 890–894.

Ayzenshtat, I., Meirovithz, E., Edelman, H., Werner-Reiss, U., Bienenstock, E., Abeles, M., and Slovin, H. (2010). Precise spatiotemporal patterns among visual cortical areas and their relation to visual stimulus processing. J Neurosci 30, 11232–11245.

Berger, T., Borgdorff, A., Crochet, S., Neubauer, F.B., Lefort, S., Fauvet, B., Ferezou, I., Carleton, A., Lüscher, H.-R., and Petersen, C.C.H. (2007). Combined voltage and calcium epifluorescence imaging in vitro and in vivo reveals subthreshold and suprathreshold dynamics of mouse barrel cortex. J. Neurophysiol. 97, 3751–3762.

Van Den Bergh, G., Zhang, B., Arckens, L., and Chino, Y.M. (2010). Receptive-field properties of V1 and V2 neurons in mice and macaque monkeys. J. Comp. Neurol. 518, 2051–2070.

Bienkowski, M.S., Mh, R., Mh, U., Benavidez, N.L., Wu, K., Gou, L., and Becerra, M. (2019). Extrastriate connectivity of the mouse dorsal lateral geniculate thalamic nucleus. J Comp Neurol 527, 1419–1442.

Bonhomme, V., Boveroux, P., Brichant, J.F., Laureys, S., and Boly, M. (2012). Neural correlates of consciousness during general anesthesia using functional magnetic resonance imaging (fMRI). Arch. Ital. Biol. 150, 155–163.

Boveroux, P., Vanhaudenhuyse, A., Bruno, M.A., Noirhomme, Q., Lauwick, S., Luxen, A., Degueldre, C., Plenevaux, A., Schnakers, C., Phillips, C., et al. (2010). Breakdown of within- and between-network resting state functional magnetic resonance imaging connectivity during propofol-induced loss of consciousness. Anesthesiology 113, 1038–1053.

Brody, C.D. (1999). Correlations without synchrony. Neural Comput. 11, 1537–1551.

Brown, R., Lau, H., and LeDoux, J.E. (2019). Understanding the Higher-Order Approach to Consciousness. Trends Cogn. Sci. 23, 754–768.

Campagna, J.A., Ph, D., Miller, K.W., Phil, D., Forman, S.A., and Ph, D. (2003). Mechanisms of Actions of Inhaled Anesthetics. N. Engl. J. Med. 348, 2110–2124.

Civillico, E.F., and Contreras, D. (2012). Spatiotemporal properties of sensory responses in vivo are strongly dependent on network context. Front. Syste 6, 1–20.

Devonshire, I.M., Grandy, T.H., Dommett, E.J., and Greenfield, S. a (2010). Effects of urethane anaesthesia on sensory processing in the rat barrel cortex revealed by combined optical imaging and electrophysiology. Eur. J. Neurosci. 32, 786–797.

Dickinson, R., White, I., Lieb, W.R., and Franks, N.P. (2000). Stereoselective Loss of Righting Reflex in Rats by. Anesthesiology 93, 837–843.

Durand, S., Iyer, R., Mizuseki, K., de Vries, S., Mihalas, S., and Reid, R.C. (2016). A Comparison of Visual Response Properties in the Lateral Geniculate Nucleus and Primary Visual Cortex of Awake and Anesthetized Mice. J. Neurosci. 36, 12144–12156.

Eger, E.I., and Johnson, B.H. (1987). Rates of awakening from anesthesia with I-653, halothane, isoflurane, and sevoflurane: a test of the effect of anesthetic concentration and duration in rats. Anesth. Analg 66, 977–982.

Eger, R.P., and MacLeod, B.A. (1995). Anaesthesia by intravenous emulsified isoflurane in mice. Can. J. Anaesth. 42:2, 173–176.

Fehérvári, T.D., and Yagi, T. (2016). Population Response Propagation to Extrastriate Areas Evoked by Intracortical Electrical Stimulation in V1. Front. Neural Circuits 10, 1–14.

Fehérvári, T.D., Okazaki, Y., Sawai, H., and Yagi, T. (2015). In Vivo Voltage-Sensitive Dye Study of Lateral Spreading of Cortical Activity in Mouse Primary Visual Cortex Induced by a Current Impulse. PLoS One 10, e0133853.

Ferezou, I., Bolea, S., and Petersen, C.C.H. (2006). Visualizing the cortical representation of whisker touch: voltage-sensitive dye imaging in freely moving mice. Neuron 50, 617–629.

Franks, N.P. (2008). General anaesthesia: from molecular targets to neuronal pathways of sleep and arousal. Nature 9, 370–386.

Gao, E., DeAngelis, G.C., and Burkhalter, A. (2010). Parallel Input Channels to Mouse Primary Visual Cortex. J. Neurosci. 30, 5921–5926.

Gao, Y., Ma, Y., Zhang, Q., Winder, A.T., and Liang, Z. (2017). Time to wake up: Studying neurovascular coupling and brain-wide circuit function in the unanesthetized animal. Neuroimage 153, 382–398.

Gilad, A., and Slovin, H. (2015). Population Responses in V1 Encode Different Figures by Response Amplitude. J. Neurosci. 35, 6335–6349.

Gilad, A., Meirovithz, E., Leshem, A., Arieli, A., and Slovin, H. (2012). Collinear Stimuli Induce Local and Cross-Areal Coherence in the Visual Cortex of Behaving Monkeys. PLoS One 7, 1–13.

Glickfeld, L.L., Reid, R.C., and Andermann, M.L. (2014). A mouse model of higher visual cortical function. Curr. Opin. Neurobiol. 24, 28–33.

Goltstein, P.M., Montijn, J.S., and Pennartz, C.M.A. (2015). Effects of Isoflurane Anesthesia on Ensemble Patterns of Ca 2 + Activity in Mouse V1: Reduced Direction Selectivity Independent of Increased Correlations in Cellular Activity. PLoS One 10(2):e011.

Greenberg, D.S., Houweling, A.R., and Kerr, J.N.D. (2008). Population imaging of ongoing neuronal activity in the visual cortex of awake rats. Nat. Neurosci. 11, 749–751.

Grinvald, A., and Hildesheim, R. (2004). VSDI: a new era in functional imaging of cortical dynamics. Nat. Rev. Neurosci. 5, 874–885.

Hentschke, H., Schwarz, C., and Antkowiak, B. (2005). Neocortex is the major target of sedative concentrations of volatile anaesthetics: strong depression of firing rates and increase of GABA A receptor-mediated inhibition. Euro 21, 93–102.

Hentschke, H., Raz, A., Krause, B.M., Murphy, C.A., and Banks, M.I. (2017). Disruption of cortical network activity by the general anaesthetic isoflurane. Br. J. Aneaesthesia 119, 685–696.

Hudetz, A.G. (2012). General Anesthesia and Human Brain Connectivity. Brain Connect. 2, 291–302.

Hudetz, A.G., and Mashour, G.A. (2016). Disconnecting Consciousness: Is There a Common Anesthetic End Point? Anesth. Analg. 123, 1228–1240.

Imas, O.A., Ropella, K.M., Ward, B.D., Wood, J.D., and Hudetz, A.G. (2005). Volatile anesthetics disrupt frontal-posterior recurrent information transfer at gamma frequencies in rat. Neurosci. Lett. 387, 145–150.

Jancke, D., Chavane, F., Naaman, S., and Grinvald, A. (2004). Imaging cortical correlates of illusion in early visual cortex. Nature 428, 423–426.

Johannesson, G., Alm, P., and Biber, B. (1984). Halothane Dissolved in Fat as an Intravenous Anaesthetic to Rats. Acta Anaesthesiol. Scand. 28, 381–384.

Kalatsky, V. a., and Stryker, M.P. (2003). New paradigm for optical imaging: Temporally encoded maps of intrinsic signal. Neuron 38, 529–545.

Katzner, S., and Weigelt, S. (2013). Visual cortical networks: Of mice and men. Curr. Opin. Neurobiol. 23, 202–206.

Krom, A.J., Marmelshtein, A., Gelbard-Sagiv, H., Tankus, A., Hayat, H., Hayat, D., Matot, I., Strauss, I., Fahoum, F., Soehle, M., et al. (2020). Anesthesia-induced loss of consciousness disrupts auditory responses beyond primary cortex. Proc. Natl. Acad. Sci. U. S. A. 117.

Lamme, V.A.F. (2006). Towards a true neural stance on consciousness. Trends Cogn. Sci. 10, 494–501.

Lau, H., and Rosenthal, D. (2011). Empirical support for higher-order theories of conscious awareness. Trends Cogn. Sci. 15, 365–373.

Liang, Z., King, J., and Zhang, N. (2012a). Intrinsic Organization of the Anesthetized Brain. J. Neurosci. 32, 10183–10191.

Liang, Z., King, J., and Zhang, N. (2012b). Anticorrelated resting-state functional connectivity in awake rat brain. Neuroimage 59, 1190–1199.

Liang, Z., Liu, X., and Zhang, N. (2015). NeuroImage Dynamic resting state functional connectivity in awake and anesthetized rodents. Neuroimage 104, 89–99.

Liu, B., Li, Y., Ma, W., Pan, C., Zhang, L.I., and Tao, H.W. (2011). Broad Inhibition Sharpens Orientation Selectivity by Expanding Input Dynamic Range in Mouse Simple Cells. Neuron 71, 542–554.

Maatuf, Y., Stern, E.A., and Slovin, H. (2016). Abnormal Population Responses in the Somatosensory Cortex of Alzheimer’s Disease Model Mice. Sci. Rep. 6, 24560.

Margalit, S.N., and Slovin, H. (2018). Spatio-temporal characteristics of population responses evoked by microstimulation in the barrel cortex. Sci. Rep. 1–14.

Marshel, J.H., Garrett, M.E., Nauhaus, I., and Callaway, E.M. (2011). Functional specialization of seven mouse visual cortical areas. Neuron 72, 1040–1054.

Masamoto, K., and Kanno, I. (2012). Anesthesia and the quantitative evaluation of neurovascular coupling. J. Cereb. Blood Flow Metab. 32, 1233–1247.

Mashour, G.A. (2005). Mechanisms of general anesthesia: from molecules to mind. Best Pract. Res. Clin. Anaesthesiol. 19, 349–364.

Mashour, G.A., Roelfsema, P., Changeux, J.P., and Dehaene, S. (2020). Conscious Processing and the Global Neuronal Workspace Hypothesis. Neuron 105, 776–798.

Michelson, N.J., and Kozai, X.T.D.Y. (2018). Isoflurane and ketamine differentially influence spontaneous and evoked laminar electrophysiology in mouse V1. J. Neurophysiol. 120, 2232–2245.

Nallasamy, N., and Tsao, D.Y. (2011). Functional Connectivity in the Brain: Effects of Anesthesia. Neurosci. 17, 94–106.

Niell, C.M., and Stryker, M.P. (2010). Modulation of Visual Responses by Behavioral State in Mouse Visual Cortex. Neuron 65, 472–479.

Oizumi, M., Albantakis, L., and Tononi, G. (2014). From the Phenomenology to the Mechanisms of Consciousness: Integrated Information Theory 3.0. PLoS Comput. Biol. 10, e1003588.

Pack, C.C., Berezovskii, V.K., and Born, R.T. (2001). Dynamic properties of neurons in cortical area MT in alert and anaesthetized macaque monkeys. Nature 414, 905–908.

Pavel, A.M., Petersen, E.N., Wang, H., Lerner, R.A., and Hansen, S.B. (2020). Studies on the mechanism of general anesthesia. PNAS 117, 13757–13766.

Petersen, C.C.H., Hahn, T.T.G., Mehta, M., Grinvald, A., and Sakmann, B. (2003). Interaction of sensory responses with spontaneous depolarization in layer 2/3 barrel cortex. PNAS 100, 13638–13643.

Pisauro, M.A., Dhruv, N.T., Carandini, M., and Benucci, A. (2013). Fast hemodynamic responses in the visual cortex of the awake mouse. J. Neurosci. 33, 18343–18351.

Polack, P.-O., and Contreras, D. (2012). Long-Range Parallel Processing and Local Recurrent Activity in the Visual Cortex of the Mouse. J. Neurosci. 32, 11120–11131.

Pullon, R.M., Lucy Yan, B., Sleigh, J.W., and E. Warnaby, C. (2020). Granger Causality of the Loss of Cortical Information Flow during Responsiveness. Anesthesiology 1–13.

Raz, A., Grady, S.M., Krause, B.M., Uhlrich, D.J., Manning, K.A., and Banks, M.I. (2014). Preferential effect of isoflurane on top-down vs. bottom-up pathways in sensory cortex. Front. Syst. Neurosci. 8, 1–22.

Redinbaugh, M.J., Phillips, J.M., Kambi, N.A., Mohanta, S., Andryk, S., Dooley, G.L., Afrasiabi, M., Raz, A., and Saalmann, Y.B. (2020). Thalamus Modulates Consciousness via Layer-Specific Control of Cortex. Neuron 106, 66–75.e12.

Rosa, M.G.P., and Krubitzer, L. a. (1999). The evolution of visual cortex: Where is V2? Trends Neurosci. 22, 242–248.

Rudolph, U., and Antkowiak, B. (2014). Molecular and neuronal substrates for general anaesthetics. Nat. Rev. 5, 709–720.

Schrouff, J., Perlbarg, V., Boly, M., Marrelec, G., Boveroux, P., Vanhaudenhuyse, A., Bruno, M., Laureys, S., Phillips, C., Pélégrini-issac, M., et al. (2011). NeuroImage Brain functional integration decreases during propofol-induced loss of consciousness. Neuroimage 57, 198–205.

Schummers, J., Yu, H., and Mriganka, S. (2008). Tuned Responses of Astrocytes and Their Influence on Hemodynamic Signals in the Visual Cortex. Science 320, 1638–1644.

Sellers, K.K., Bennett, D. V, Hutt, A., Williams, J.H., and Fröhlich, F. (2015). Awake vs. anesthetized: layer-specific sensory processing in visual cortex and functional connectivity between cortical areas. J. Neurophysiol. 3798–3815.

Shushruth, S. (2013). Exploring the Neural Basis of Consciousness through Anesthesia. J. Neurosci. 33, 1757–1758.

Sonner, J.M., Werner, D.F., Elsen, F.P., Ph, D., Xing, Y., Liao, M., Harris, R.A., Ph, D., Harrison, N.L., Ph, D., et al. (2007). Effect of isoflurane and other potent inhaled anesthetics on minimum alveolar concentration, learning, and the righting reflex in mice engineered to express α1 γ-aminobutyric acid type A receptors unresponsive to isoflurane. Anesthesiology 106, 107–113.

Stettler, D.D., Das, A., Bennett, J., and Gilbert, C.D. (2002). Lateral connectivity and contextual interactions in macaque primary visual cortex. Neuron 36, 739–750.

Vaiceliunaite, A., Erisken, S., Franzen, F., Katzner, S., and Busse, L. (2013). Spatial integration in mouse primary visual cortex. J. Neurophysiol. 110, 964–972.

Wahrenbrock, E.A., Eger, E.I., Laravuso, R.B., and Maruschak, G. (1974). Anesthetic uptake - of mice and men (and whales). Anesthesiology 40, 19–2.

Wang, Q., and Burkhalter, A. (2007). Area map of mouse visual cortex. J. Comp. Neurol. 504, 287–297.

Wang, Q., Gao, E., and Burkhalter, A. (2011). Gateways of ventral and dorsal streams in mouse visual cortex. J. Neurosci. 31, 1905–1918.

Wang, Q., Sporns, O., and Burkhalter, a. (2012). Network Analysis of Corticocortical Connections Reveals Ventral and Dorsal Processing Streams in Mouse Visual Cortex. J. Neurosci. 32, 4386–4399.

White, N.S., and Alkire, M.T. (2003). Impaired thalamocortical connectivity in humans during general-anesthetic-induced unconsciousness. Neuroimage 19, 402–411.

Wollstadt, P., Sellers, K.K., Rudelt, L., and Priesemann, V. (2017). Breakdown of local information processing may underlie isoflurane anesthesia effects. Plos Comput. Biol. 13, e1005511.

Wu, T.L., Mishra, A., Wang, F., Yang, P.F., Gore, J.C., and Chen, L.M. (2016). Effects of isoflurane anesthesia on resting-state fMRI signals and functional connectivity within primary somatosensory cortex of monkeys. Brain Behav. 6, 1–12.

Yang, W., Chini, M., Pöpplau, J.A., Formozov, A., Piechocinski, P., Rais, C., Morellini, F., Sporns, O., Hanganu-Opatz, I.L., and Wiegert, J.S. (2020). Anesthetics uniquely decorrelate hippocampal network activity, alter spine dynamics and affect memory consolidation. BioRxiv 2020.06.05.135905.

